# The Effect of Kinship in Re-identification Attacks Against Genomic Data Sharing Beacons

**DOI:** 10.1101/2020.01.30.926907

**Authors:** Kerem Ayoz, Miray Aysen, Erman Ayday, A. Ercument Cicek

**Affiliations:** Computer Engineering Department, Bilkent University, Ankara, 06800, Turkey; Computer and Data Sciences Department, Case Western Reserve University, Cleveland, OH 44106; Computational Biology Department, Carnegie Mellon University, Pittsburgh, PA 15213

## Abstract

**Motivation:** Big data era in genomics promises a breakthrough in medicine, but sharing data in a private manner limits the pace of field. Widely accepted “genomic data-sharing beacon” protocol provides a standardized and secure interface for querying the genomic datasets. The data is only shared if the desired information (e.g., a certain variant) exists in the dataset. Various studies showed that beacons are vulnerable to re-identification (or membership inference) attacks. As beacons are generally associated with sensitive phenotype information, re-identification creates a significant risk for the participants. Unfortunately, proposed countermeasures against such attacks have failed to be effective, as they do not consider the utility of beacon protocol.

**Results:** In this study, for the first time, we analyze the mitigation effect of the kinship relationships among beacon participants against re-identification attacks. We argue that having multiple family members in a beacon can garble the information for attacks since a substantial number of variants are shared among kin-related people. Using family genomes from HapMap and synthetically-generated datasets, we show that having one of the parents of a victim in the beacon causes (i) significant decrease in the power of attacks and (ii) substantial increase in the number of queries needed to confirm an individual’s beacon membership. We also show how the protection effect attenuates when more distant relatives, such as grandparents are included alongside the victim. Furthermore, we quantify the utility loss due adding relatives and show that it is smaller compared to flipping based techniques.

## 1 Introduction

In the last two decades, emerging sequencing technologies have been providing researchers with larger genomic datasets which creates new opportunities for understanding the genetic architectures of diseases and have been providing insights for new therapies (Kim, 2001). This was further fueled by the exponential growth of the personal genomics industry in the last five years which attracted consumers that want to (i) familiarize themselves with their genetic origins or (ii) take precautions against possible health risks (Khan and Mittelman, 2018). Growing size of genomic datasets promises new opportunities for research through data sharing. However, data inherently contains highly sensitive information and privacy-preserving and secure sharing of data comes up as a major challenge. Anonymization of the genomes is a straightforward solution. However, the genome is the utmost personal identifier and it can reveal the identity of an individual. Such a scenario can dire ethical consequences, such as discrimination (e.g., on the basis of employment or insurance (Kim, 2001; Billings *et al*., 1992; Lapham *et al*., 1996)). Leakage of genomic information of an individual not only jeopardises their privacy but also the privacy of their relatives since genomic information of an individual can be used to infer genomes (and hence genetic predisposition to a diseases) of other family members (Humbert *et al*., 2013). For instance, Deznabi *et al*., demonstrate that the single nucleotide polymorphisms (SNPs) of relatives can be reconstructed with high confidence using (i) Mendel’s law, (ii) high-order correlations between SNPs, and (iii) minor allele frequencies (MAFs) of the SNPs in a population. Thus, researchers face a trade-off between (i) sharing data to empower genetic research, which puts the participants under risk and legally binds them for possible repercussion and (ii) not sharing the data, which potentially bars the advances in life sciences.

In 2016, the Global Alliance for Genomics and Health (GA4GH) introduced the Beacon Project, a system constructed with the aim of providing a secure and systematic way of sharing genomic data. Beacons provide an interface, in which a user can query the existence of a specific nucleotide at a given position in a particular chromosome. For instance, “is there a participant carrying nucleotide C at the 100,000^*th*^ position of chromosome 1?” is a valid query. The beacon responds to such a query only with a simple “yes” or “no”. Therefore, the beacon protocol is considered safer (compared to other statistical databases), as the query responses are binary and they do not include any information about allele frequencies. Moreover, a “yes” answer cannot be tied to a specific individual in the beacon. The beacon protocol also encourages cross site collaborations because the users do not have to go through the rigorous paperwork unless they identify a useful dataset for their research.

Nonetheless, previous studies showed that beacons are vulnerable to reidentification attacks (Shringarpure and Bustamante, 2015; Raisaro *et al*., 2017; von Thenen *et al*., 2018). Shringarpure and Bustamante showed that a likelihood ratio test (LRT) can be used to infer the membership of an individual to a beacon by querying the beacon for a couple of hundred SNPs of that individual (SB Attack). This study clearly showed that the beacons indeed leak information which potentially leads to the disclosure of sensitive information if the beacon is associated with a sensitive trait (e.g., SFARI beacon which contains participants with autism). Raisaro *et al*. advanced the SB Attack by assuming the attacker has information regarding the MAFs in the population (Optimal Attack). By asking SNPs with low MAF values first, they showed that an attacker actually needs only a handful of queries to achieve the same power as the SB Attack. Finally, von Thenen *et al*. introduced two new attacks. First, they showed that the attacker can infer beacon responses using the responses of previously asked queries (Query Inference - QI Attack). Second, they showed that the attacker can still launch an attack even if the victim has concealed their SNPs with low MAF values (Genome Inference - GI Attack). Both attacks utilize the correlations among SNPs and they further decrease the number of required queries for confident inference.

Several countermeasures have been proposed in the literature to protect the privacy of the beacon participants against re-identification attacks. Shringarpure and Bustamante considered: (i) having larger beacon sizes, (ii) sharing only small genomic regions (e.g., genes of interest) instead of full genome, (iii) having a uniform ancestry composition in the beacon, and (iv) not publishing the metadata (e.g., dataset size). However, as also stated by the authors, these techniques reduce the utility of the beacon. Raisaro *et al*. proposed a query budget per participant, which expires if many SNPs of an individual is queried and the participant is taken out of the system (i.e., queries including them are not answered). Yet, von Thenen *et al*. showed that inference of beacon answers via SNP correlations can get around such budget-based countermeasures. Al Aziz *et al*. proposed two algorithms that randomize the beacon responses. However, such noise-based techniques reduce the utility of the users and substantially affect the usability of the system.

Another line of work propose randomly flipping beacon responses to reduce the power of re-identification attacks. Bu *et al*. showed that flipping a certain amount of rare SNPs in the beacon responses can reduce the re-identification power to an insignificant level. However, flipping the responses to the queries that are received for rare SNPs is shown to significantly reduce the utility of beacon responses. Thus, Bu *et al*. proposed a real-time flipping (RTF) method which aims at flipping the queries that are received for the rare SNPs of more vulnerable individuals in the beacon. The difference between RTF and other flipping methods is that it guarantees the same level of privacy by flipping fewer responses. RTF method achieves this goal by using a *p*-value for each potentially target individual in the beacon. *p* value of a potential target is the fraction of LRT scores in a randomly selected control group that is equal to or smaller than that of the target individual. If the *p*-value is any of an individual in the beacon is smaller than 5%, providing the correct response of the query is assumed to significantly increase the vulnerability of the corresponding individual for the re-identification attack. Thus, in that case, RTF flips the response of the query. Although RTF method performs better than other flipping methods (in terms of utility), it significantly reduces the utility for the beacon responses for rare SNPs.

In this study, we consider using the kinship of beacon participants as a countermeasure against re-identification attacks. We show that the power of the state-of-the-art attacks substantially decrease when at least one of the parents of a victim is added to the beacon. The key idea is that kinship garbles the information returned to the attacker since family members share many SNPs and the re-identification attack algorithm cannot conclude weather the “yes” answer coming from the beacon originates due to the victim or their relatives.

Using a beacon constructed from the CEU population of the HapMap dataset, we show that the number of queries to infer beacon membership of a victim increase when at least one of their family members is added to the beacon. We also show how the power loss for the state-of-the-art re-identification attacks changes with different degrees of relatives in the beacon. Finally, we quantify the utility loss of the beacon due to this proposed mitigation technique. We define the utility as the proportion of the flipped beacon responses (due to the proposed mitigation technique) and we show that the proposed mitigation technique does not cause a significant decrease in utility (especially for SNPs with low MAF values).

The rest of the paper is organized as follows. In Section 2, we provide technical details on methods and datasets we use. In Sections 3 and 4, we provide and discuss the results about the effect of kinship on reidentification attacks under various settings. Finally, in Section 5, we conclude the paper.

## 2 Materials and Methodology

In this section, we present the technical details on the state-of-the-art re-identification attacks against genomic data sharing beacons. We also describe our techniques to quantify the power of the attacker and the family simulation procedure (for the evaluations we conduct using synthetic datasets).

### 2.1 Re-identification Attacks Against Beacons

Shringarpure and Bustamante introduced the first re-identification attack against beacons. The algorithm repeatedly queries for a victim’s heterozygous SNPs and a likelihood ratio test (LRT) is performed to choose between a null hypothesis (*H*_0_, in which the queried genome is not in the beacon) and an alternative hypothesis (*H*_1_, in which the queried genome is a member of the beacon). The log-likelihood (*L*) under the null and alternate hypothesis are shown as follows:

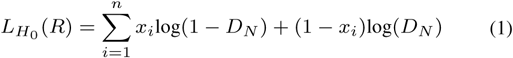

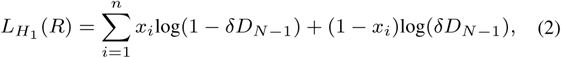

where *R* is the set of responses and *x*_*i*_ is the binary response for the *i*^*th*^ query (*x*_*i*_ = 1 if the query response is “yes” and *x*_*i*_ = 0, otherwise). *δ* term in the alternate hypothesis indicates a small probability that attacker’s copy of the victim’s genome does not match the beacon’s copy (e.g., due to differences in the sequencing pipelines). *n* is the number of queried SNPs, *D*_*N*_ is the probability that none of the *N* individuals in the beacon has the corresponding allele for the queried SNP, and *D*_*N*−1_ is the probability that no individual except for the victim having the corresponding allele for the queried SNP. The LRT statistic Λ is calculated as follows:

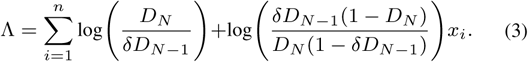

*H*_0_ is rejected if it is less than a threshold and this threshold can be found theoretically under the assumption that the queried SNPs are i.i.d. Raisaro *et al*., introduced the Optimal Attack, which assumes that the attacker has access to the minor allele frequencies (MAF) of a population representing the beacon participants. Then, the SNPs are queried in the ascending MAF order. The formulation is identical to the SB Attack, but in Optimal Attack, the computations of *D*_*N*−1_ and *D*_*N*_ depend on the query *i* since each query has a different effect on the LRT statistic. Thus, in Optimal Attack, 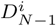 and 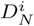 are calculated as follows: 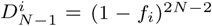 and 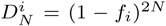, where *f*_*i*_ represents the MAF of SNP *i*. The likelihood-ratio test (LRT) statistic, Λ is then computed as follows:

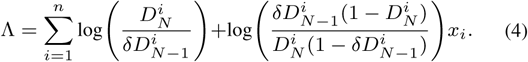

The Λ threshold (*t* _*α*_) for rejection of the null hypothesis is determined empirically for every query since the i.i.d. assumption in Shringarpure and Bustamante (2015) no longer holds. That is, for every query, the distribution of Λ under the null hypothesis is found using *k* individuals that are not in the beacon. When Λ value of a victim is less than *tα*, the alternative hypothesis is chosen, where *α* represents the false positive rate.

von Thenen *et al*. (2018) introduced the Query Inference (QI) Attack, which extends the Optimal Attack by showing that in addition to the MAF information of population, the attacker can also utilize the correlations between the SNPs. The correlation between the SNPs are calculated based on their LD values and a SNP-SNP network is generated, in which the vertices are the SNPs and the weights on directed edges represent the LD values. When a SNP is queried, the beacon responses of the neighboring SNPs in the SNP-SNP network are inferred, and hence the required number of queries is significantly decreased. In the QI Attack, the null and the alternative hypothesis formulations (and the corresponding Λ definition) are changed, so that the new calculation also reflects the information obtained from (*m*) inferred queries. The inferred queries are weighted by an inference confidence (*γ*) and the new log-likelihoods and LRT statistics are computed as follows:

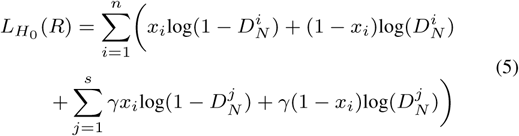

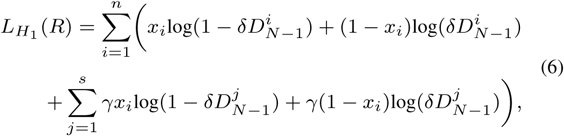

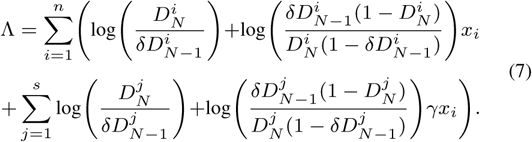

### 2.2 Power Calculation

We perform a power analysis to quantify the success of a re-identification attack (Raisaro *et al*., 2017; von Thenen *et al*., 2018). All Optimal, QI and GI Attacks query SNPs in the ascending MAF order. In this scheme, for every query *i*, a 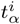 value is determined which is the Λ threshold to reject the NULL hypothesis. *α* represents the desired false positive rate. We pick *k* people (controls) who are not in the beacon. We assume these *k* people has a similar population structure as the beacon participants. For each of the *k* controls, the *i*^*th*^ query is posed and a Λ value set is obtained: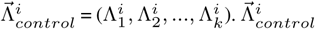 is sorted (in ascending order) and the Λ^*i*^ value, for which *α* percent of *k* people have smaller Λ^*i*^s, is picked as the 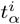. For instance, if *k* is 20 and *α* is 0.05, then the second smallest Λ^*i*^ is picked as 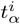 as 1 person is below that threshold. This represents the false positive threshold as for that person, the NULL hypothesis would have been rejected. Given *n* queries, the 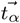 list is generated which contains the *tα* values of all *n* queries: 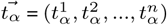.

To measure the power per each query *i*, first, *l* people from the beacon (cases) are obtained. Then, for everyone in this set, the *i*^*th*^ query is posed and a Λ value set is obtained: 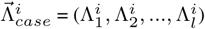. Then, the power for the *i*^*th*^ query is calculated as follows: 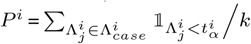. This is the fraction of the cases who have 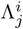 value that is less than 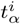. For that fraction of the *l* people, the NULL hypothesis is (correctly) rejected. For instance, if *l* is 20 and 5 people have 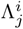 that is less than 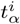, this means for the *i*^*th*^ query, the power of the attack is 25% at the *α* false positive rate. The vector 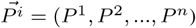 is then plotted to see the power change with respect to increasing number of queries. An attack which reaches to 100% power earlier than others is considered more powerful. Our goal in using kinship as a countermeasure aims at decreasing the power of the state-of-the-art attacks, and thus either increasing the number of queries to reach 100% power or preventing the attack to reach 100% power at all.

### 2.3 Generation of Relatives’ Genomes

The family genome data we use for our experiments (which is obtained from HapMap as discussed in Section 3.1) contains only trios (mother, father and the child). Thus, we also generate synthetic parent genomes for a given victim’s (child’s) actual genome to be able to simulate the effect of having more distant relatives in the beacon such as grandparents.

Flow of the relative generation algorithm is shown in Figure 1. When creating parents’ genomes, in order to preserve haploblock structure, we separated child’s alleles to two different strands in a given block which has a size of 18kb, which is the average block size for human genome Consortium *et al*. (2005). Since we did not have strand information for minor allele in heterozygous SNPs (i.e., phasing), we distributed minor alleles to strands randomly in heterozygous SNPs. As a result, we obtained single strands for both father and mother. Generation of the remaining strand for both parents are same. For each allele in remaining strand, we picked either a major allele or a minor allele according to the allele frequency of each that SNP. After creating both strands of the parents, we obtained SNPs of the parents by joining strands together. We used the same algorithm to generate the genomes of the victim’s grandparents, using the generated parents as the child. We also assume that generated couples (i.e., parents and grandparents) are not related. Table 1 shows the possible genotypes of the parents given the genotype of the child.

**Table 1:**
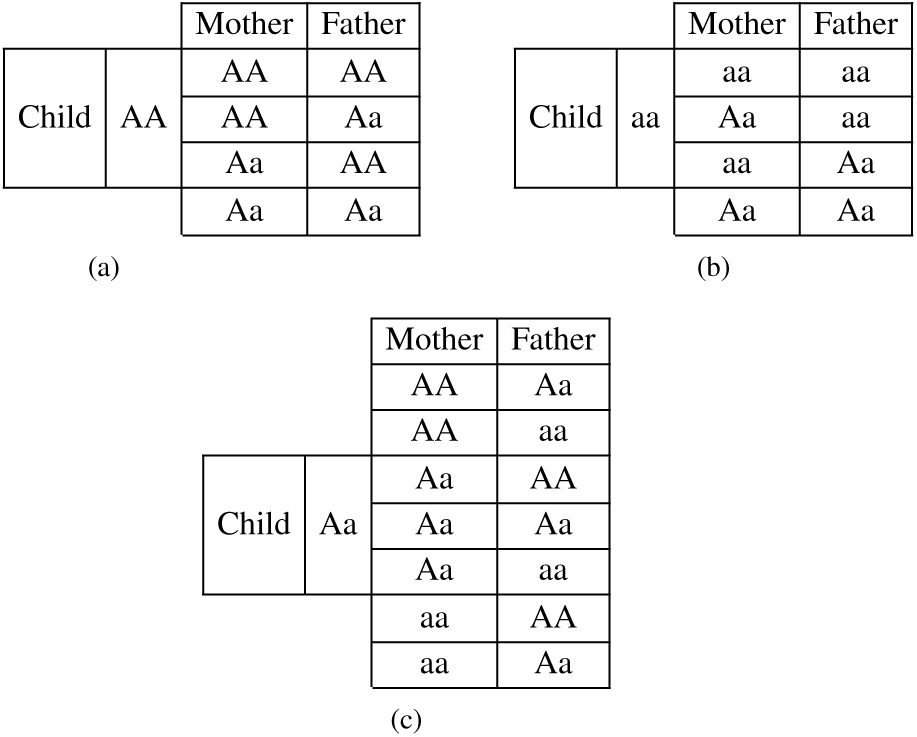
A toy example showing the possible SNPs of the parents for three cases, in which the child’s SNP is (a) major homozygous, (b) minor homozygous, and (c) heterozygous.

**Fig. 1:**
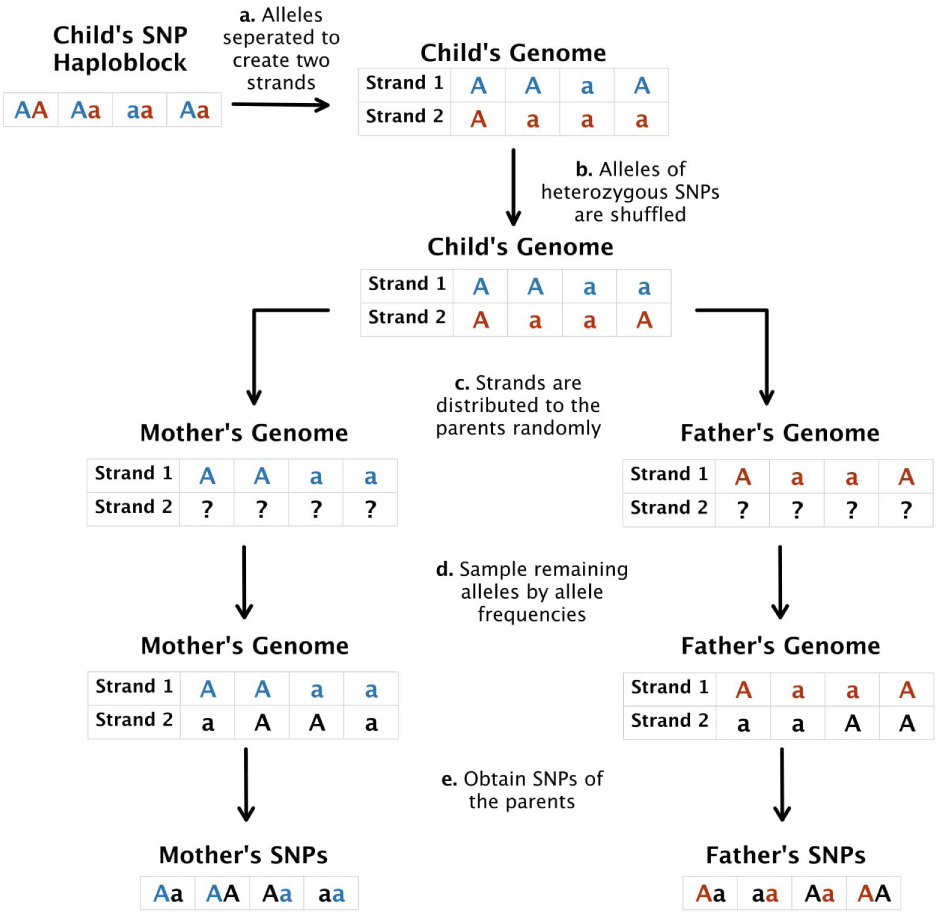
Flow of family member generation algorithm. First, a haploblock for the child (i.e., the victim) is obtained. (a) Two strands of the DNA for the child is formed by separating the alleles. (b) The minor alleles are randomly shuffled to form the final version of each strand (blue for strand 1 and red for strand 2). (c) These strands are used to create the first strand of each parent. (d) The second strand for each parent is constructed by randomly assigning the minor allele while taking the minor allele frequency into account. (e) Genotypes of the parents are obtained for the corresponding haploblock.

## 3 Results

In this section, we provide the results showing the decrease in the attacker’s power once a relative (or a set of relatives) is added to the beacon. We also quantify the utility loss in beacon responses once a relative of the victim is added to the beacon for privacy protection.

### 3.1 Experimental Setup

For our experiments, we obtained the genomes of the Utah residents with northern and western European ancestry (CEU) population of the HapMap project (Consortium *et al*., 2003). The same population and similar beacon sizes are also used in all previous re-identification attacks (Shringarpure and Bustamante, 2015; Raisaro *et al*., 2017; von Thenen *et al*., 2018). For our experiments with synthetically generated families, we created artificial genomes for the parents and grandparents of selected victim(s) from this population as described in Section 2.3. The original dataset also contains 40 real families (i.e., trios) and we used this data to show the results on actual genomes of the family members. Note that the original dataset does not contain actual genomes of grandparents. We calculated the MAF values (needed for the Optimal and QI Attacks) from the HapMap dataset using 100 individuals from this population. The QI Attack requires the LD values for the considered SNPs to create the SNP-SNP network for query inference. We used the same 100 individuals from the HapMap dataset to create these models.

To test the effect of having family members in the beacon along with a victim, we compare two cases: (i) the victim is in the beacon and no other relatives are, and (ii) the victim is in the beacon and one or more relatives are in the beacon with her. Thus, we have two power calculation settings for each of these cases. Case (i) is straightforward and performed as also done in Raisaro *et al*. (2017); von Thenen *et al*. (2018) and as detailed in Section 2.2. Case (ii) requires the beacon to contain the family members of the victim and it requires the following adjustments: First, 20 individuals are selected from the CEU population of the HapMap dataset. When these 20 people are used as controls, by definition, they are excluded from the beacon, but their considered relative(s) for that test (e.g., 20 mothers) are in the beacon along with 40 unrelated individuals from the same population. After determining the 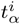 for every *i*^*th*^ query, these 20 people are now considered as the cases. Now, the beacon contains these 20 people and their relative(s) along with 20 unrelated people. For the tests with synthetically generated data, synthetic parents and grandparents of 20 CEU individuals are generated as described in Section 2.3 and the above procedure is performed similarly.

### 3.2 Re-identification Attacks on Genomic Data Sharing Beacons with Family Members

In this study, we argue that adding family members to the beacon will improve the privacy of beacon participants, and hence it will be a natural mitigation technique. The origin of this idea is the inheritance, the fact that an individual’s genome is constructed based on their parents genomic information. We also claim that addition of family members to the beacon does not cause a significant utility loss. We further discuss this in Section 3.3. To show how the results of attacks change with the presence of family members in the beacon, we used the same experiment parameters as the previous re-identification attacks (detailed information about datasets and experimental settings are in Section 3.1).

The attacker’s goal is to infer whether the targeted individual (victim) is in the beacon or not. The attacker has the following auxiliary information along with the VCF of the victim: MAF of the victim’s population and LD values. We let *t* be an evaluation parameter representing the threshold for the hidden SNPs of the victim (e.g., as a countermeasure against the re-identification attack). That is, we assume the victim hides their SNPs with MAF values less than *t*. We let the attacker query the beacon for the heterozygous SNP positions of the victim (to have the same settings with previous re-identification attacks).

We assume that the attacker does not have access to VCF files of victim’s family members. The attacker may or may not know the existence of victim’s family members in the beacon since this knowledge does not provide an advantage to the attacker to infer the membership of the victim. If attacker knows that (at least) a family member is in the beacon, it cannot be sure about the reason of the “yes” responses (e.g., whether they are due to the victim or other family members). If the attacker does not know the membership information of victim’s family members, it will possibly come to a wrong conclusion about the membership of the victim to the beacon. Thus, in both cases, attacker’s inference power for the victim’s membership will be low (due to the existence of family members in the beacon).

We performed the Optimal and QI Attacks for different scenarios for the individuals in the beacon: (i) the original beacon that does not involve victim’s family members, (ii) beacon that contains victim’s mother, (iii) beacon that contains victim’s father, (iv) beacon that contains victim’s both parents, (v) beacon that contains victim’s grandparents from mother’s side (only for synthetically generated genomes), and (vi) beacon that contains victim’s grandparents from father’s side (only for synthetically generated genomes). We show these different scenarios in Figure 2.

**Fig. 2:**
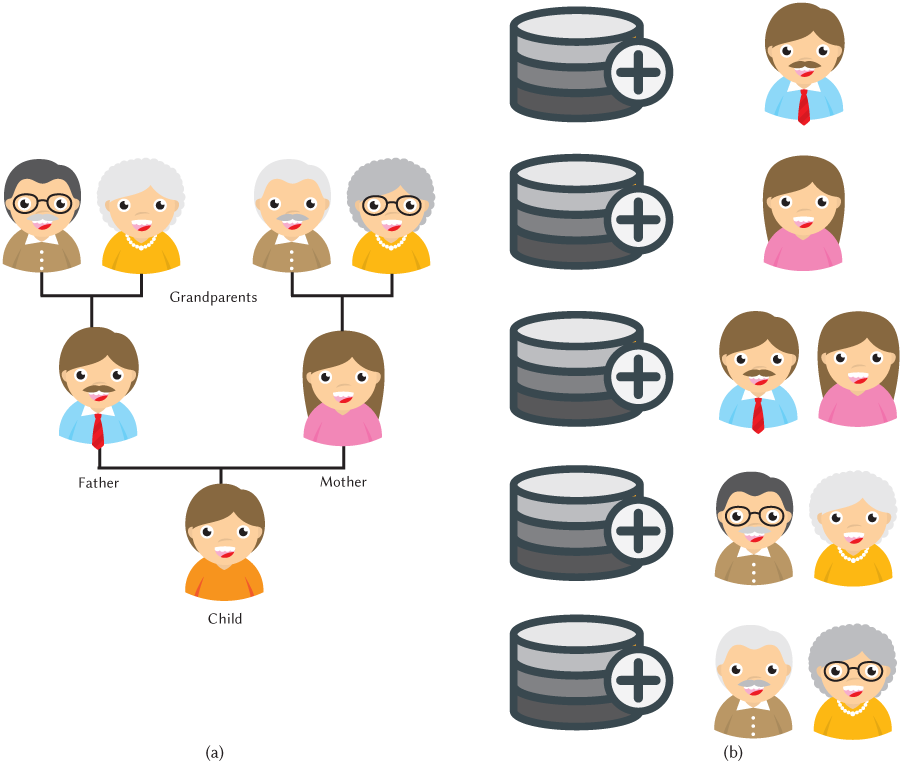
(a) Family tree of the victim (child in the figure). (b) Experimental setup for different scenarios for the individuals in the beacon.

First, we show how the power of the attack changes for (i) the original beacon (that does not include any family members of the victim), (ii) the beacon that only includes the mother of the victim, (iii) the beacon that only includes the father of the victim, and (iv) the beacon that includes both parents of the victim. Figure 2 shows the settings we consider.

Figure 3 shows the results obtained with the synthetic parents and Figure 4 shows the results obtained with the actual parents. We observed that both experiments follow the same pattern while the power loss in the experiments with synthetic data is slightly more. In von Thenen *et al*. (2018), authors show that the individual’s membership to the beacon can be inferred with high power with only a few queries. Our results for the original beacon (that does not include any family members of the victim) are also consistent with the results of von Thenen *et al*. (2018). We also observed that when at least one family member of the victim is in the beacon, the power curves shift to right, meaning that the attacker needs more queries to infer the membership of the victim to the beacon. For instance, when at least one family member of the victim is in the beacon, the power only reaches to 0.1 after two queries (for which the QI Attack’s power reaches to 1 for the original beacon).

**Fig. 3:**
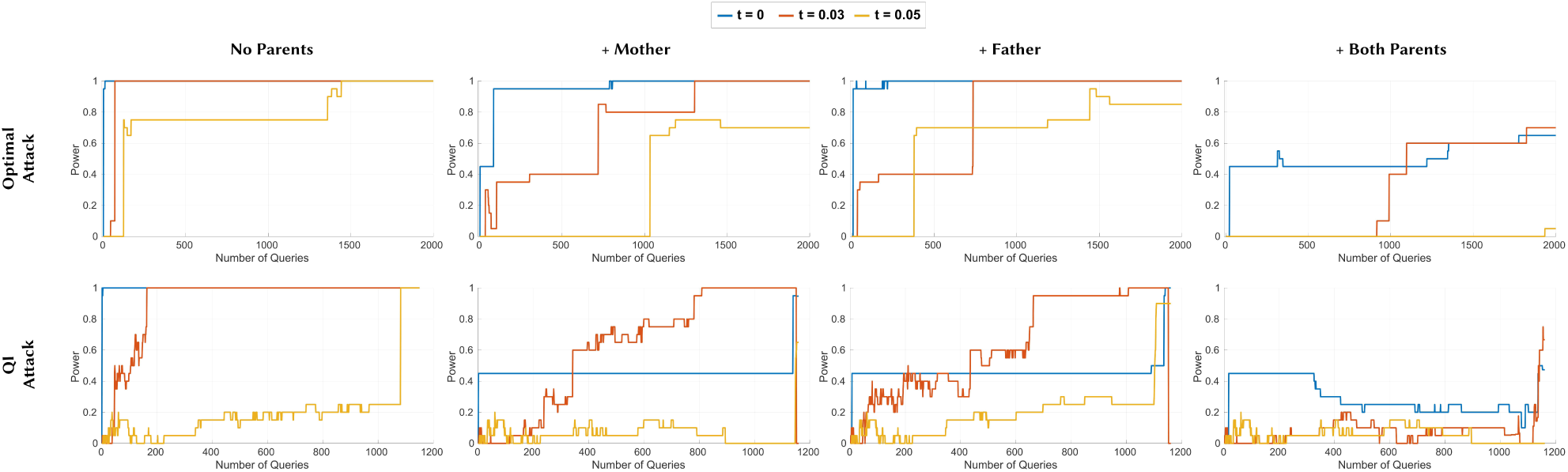
The power curves of the Optimal Attack and QI Attack with different beacon setups with synthetic family members. For all attacks: (i) the first plot is when the beacon does not include any family members of the victim, (ii) the second plot is when only the mother of the victim is in the beacon, (iii) the third plot is when only the father of the victim is in the beacon, and (iv) the fourth plot is when both parents of the victim are in the beacon. SNPs of the victim with MAF values smaller than *t* are hidden from the attacker.

**Fig. 4:**
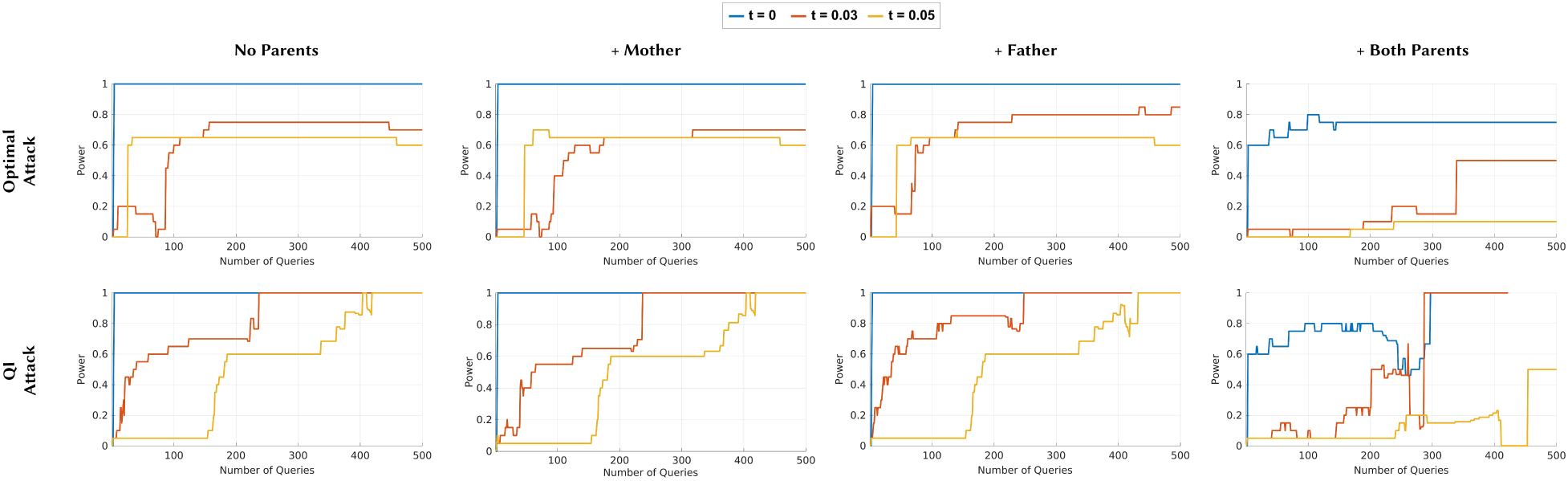
The power curves of the Optimal Attack and QI Attack with different beacon setups with actual family members. For all attacks: (i) the first plot is when the beacon does not include any family members of the victim, (ii) the second plot is when only the mother of the victim is in the beacon, (iii) the third plot is when only the father of the victim is in the beacon, and (iv) the fourth plot is when both parents of the victim are in the beacon. SNPs of the victim with MAF values smaller than *t* are hidden from the attacker.

We also observed that in the Optimal Attack, when *t* = 0, including only the mother or father of the victim to the beacon increases the number of queries for the attacker (to have a high power) to hundreds. In the QI Attack on the other hand, we observed that when at least one family member of the victim is in the beacon, the attacker’s power reaches to 1 in hundreds of queries for only smaller values of *t*. Furthermore, for all attacks, when both parents of the victim are in the beacon, attacker’s power never reaches to 1, and it is always low. This is expected since the minor alleles of the child (victim) either come from the mother or the father. Thus, when both parents of the victim are in the beacon, there is no way for the attacker to make inference about the membership of the victim. We also observed that once the power converges to a value, it does not change even if the attacker keeps asking for more queries.

We have shown in the above experiment that the synthetically generated genomes of the victim’s parents provide highly correlated and less optimistic results compared to the experiment with actual parents’ genomes. Relying on this fact, we used synthetic genomes of the grandparents to simulate the effect of the existence of more distant relatives in the beacon. That is, we showed how adding grandparents to the beacon affect the attacker’s power for the re-identification using the Optimal Attack. As we show in Figure 5, adding only one of the grandparents to the beacon (mother’s father as in Figure 5(a) or mother’s mother as in Figure 5(b)) causes the attacker’s power decrease less than adding the mother (in Figure 3) since degree of kinship decreases. In other words, as expected, the decrease in attacker’s power is inversely proportional with the distance between the victim and their relatives. We also obtained similar results when we added father’s mother and father’s mother separately. Furthermore, we observe that adding mother’s both parents (i.e., victims grandparents from mother’s side as in Figure 5(c)) to the beacon is almost equivalent to adding the mother. Similarly, adding father’s parents to the beacon (as in Figure 5(d)) is almost equivalent to adding the father. Note however that adding mother’s (or father’s) both parents provide a slightly stronger mitigation compared to adding only the mother (or father). This is because adding mother’s both parents introduce more diversity to the beacon compared to adding just the mother. For instance, comparing the beacon including only victim’s mother and mother’s parents, the beacon including victim’s grandparents may include more “yes” responses (due to heterozygous SNPs of the grandparents that may not occur in the mother).

**Fig. 5:**
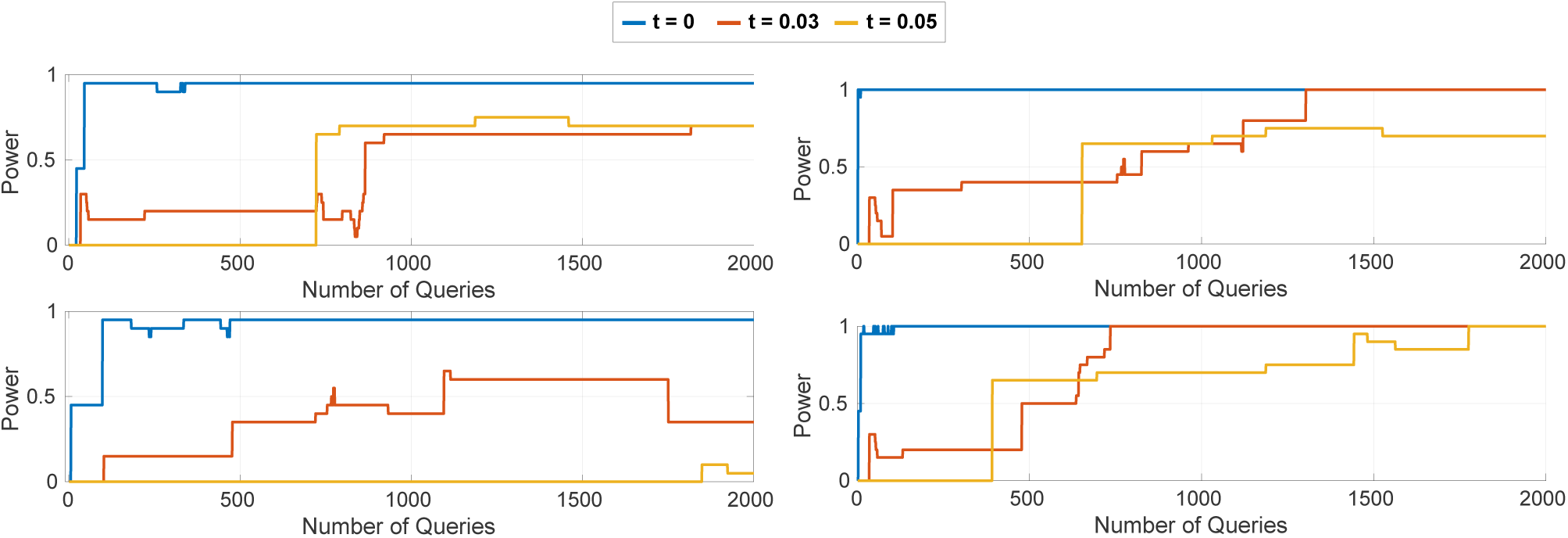
The power curves of the Optimal Attack when the beacon includes (top-left) victim’s mother’s father, (top-right) victim’s mother’s mother, (bottom-left) victim’s both grandparents from mother’s side, and (bottom-right) victim’s both grandparents from father’s side.

### 3.3 Utility Analysis of the Proposed Mitigation Technique

We showed that adding victim’s family members to the beacon significantly increases the number of queries needed for the attacker to have a high power. However, as discussed, beacons are typically associated with a particular phenotype (i.e., all participants of the beacon has the corresponding phenotype). Thus, adding a family member of the victim to the beacon may result in a utility decrease in beacon’s responses since (i) the added family member(s) may not have the corresponding phenotype of the beacon and (ii) the added family member(s) may result in a change in beacon’s original responses.

In particular, if the original beacon response (before adding any family members as a mitigation technique) is “no” for a query and adding a family member changes that beacon response to “yes” (due to heterozygous SNPs of the added family member), utility of beacon decreases. Therefore, we define the utility loss of the beacon as the fraction of additional “yes” responses that arise due to the addition of one or more extra individuals (family members of the victim) as a result of the proposed mitigation technique. In Tables 2 and 3, we show the decrease in utility of beacon’s responses for both case and control groups (that we used in our experiments) due to the addition of the family member(s) as a mitigation technique. Note that corresponding family members are added together at the same time for all 20 cases and 20 controls, respectively. That is, we observe that the utility loss is less than 10% even when both parents of the victim are included in the beacon using the synthetic dataset and, less than 8% using the actual family members.

**Table 2:** The fraction of additional “yes” responses that arise due to the addition of family members (synthetic) of the victim (as a result of the proposed mitigation technique) is shown as a measure of utility loss for the case and control groups. Each individual in the case and control groups are selected as the victim and are added to the CEU beacon with 65 individuals (note that cases are already in the beacon). The utility loss is calculated when all parents are added to the beacon at the same time.

|  | Mother in Beacon | Father in Beacon | Both Parents in Beacon |
| --- | --- | --- | --- |
| Control Group | 4.50% | 7.52% | 9.78% |
| Case Group | 2.67% | 6.78% | 9.13% |

**Table 3:** The fraction of additional “yes” responses that arise due to the addition of family members (actual) of the victim (as a result of the proposed mitigation technique) is shown as a measure of utility loss for the case and control groups. Each individual in the case and control groups are selected as the victim and are added to the CEU beacon with 60 individuals (note that cases are already in the beacon). The utility loss is calculated when all parents are added to the beacon at the same time.

|  | Mother in Beacon | Father in Beacon | Both Parents in Beacon |
| --- | --- | --- | --- |
| Control Group | 5.05% | 4.89% | 7.96% |
| Case Group | 3.24% | 3.36% | 5.75% |

SNPs with a lower MAF values are particularly important for the researchers since there is an inverse relationship between the a variant’s disease odds ratio and its frequency (Bomba *et al*., 2017). Thus, we also quantified the utility loss of the beacon responses considering the SNPs with low MAF values. One by one, we added the mothers of 20 case individuals to the beacon and observed the utility loss for various MAF thresholds. For the synthetically generated mothers, Figure 6 shows the utility loss in beacon responses (y-axis) for all SNPs with an MAF value less than a threshold (x-axis; cumulative). We observed that utility loss is substantially smaller for SNPs with lower MAF values. For instance, for SNPs with an MAF less than 0.01, the utility loss is less than 0.05%. We observed a similar trend in Figure 7, which shows the results when actual mothers are added to the beacon in the same manner. Similar to before, we observed that the results obtained in the synthetic genomes is overly conservative and when actual mothers are used, the utility loss is roughly 4 folds less, which shows that the utility is mostly preserved despite a substantial power loss of the attacker as shown in Figure 4. In other words, adding a family member of the victim to the beacon does not cause much change in the results of queries that involve low-MAF SNPs, which is expected as such SNPs are rare and are not frequently observed.

**Fig. 6:**
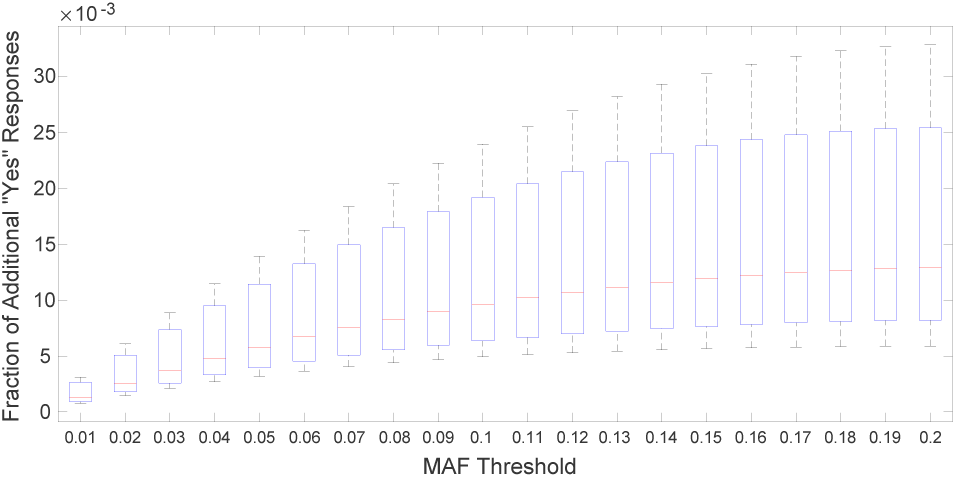
The utility loss of the beacon responses considering the SNPs with low MAF values. The box plots show the fraction of additional “yes” responses that arise due to the addition of synthetically generated family members for 20 cases when the mother of each victim is added to the beacon one-by-one independently. The x-axis shows various MAF thresholds. For each x value, all SNPs with an MAF value less than that threshold are considered.

**Fig. 7:**
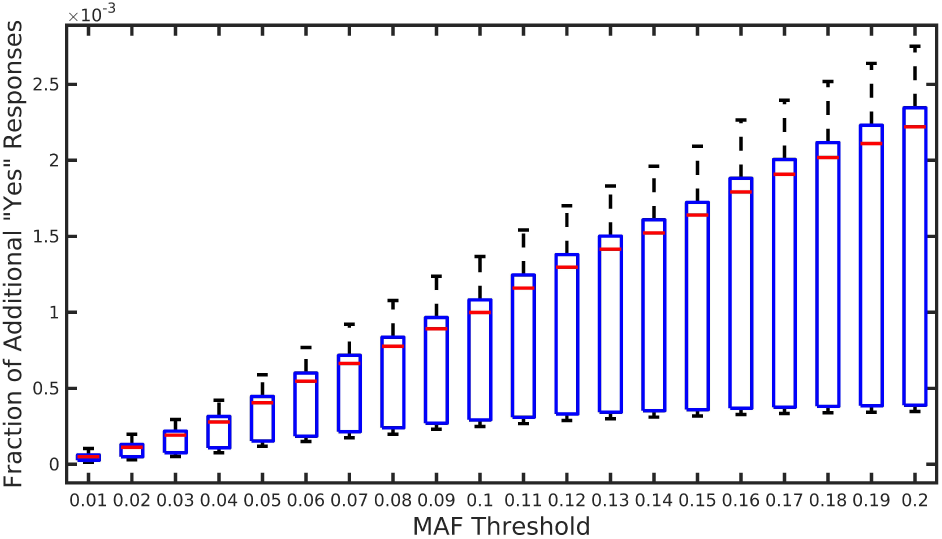
The utility loss of the beacon responses considering the SNPs with low MAF values. The box plots show the fraction of additional “yes” responses that arise due to the addition of actual family members for 20 cases when the mother of each victim is added to the beacon one-by-one independently. The x-axis shows various MAF thresholds. For each x value, all SNPs with an MAF value less than that threshold are considered.

## 4 Discussion

Genomic data sharing beacon protocol has been widely accepted by the community as the golden standard for secure and privacy preserving data sharing. The Beacon Network^1^, providing a central querying mechanism to 80 beacons, lit all over the world for various phenotypes ranging from autism to cancer (accessed on January 28, 2020). However, the information leaks identified by several re-identification attack algorithms and by a recently introduced genome reconstruction attack (Ayoz *et al*., 2020) questions the usability of the system. Currently, setting up a beacon is a risk for all parties including genome donors, data owners, and even for the beacon system operators due to possible ethical, legal, and monetary repercussions.

A correspondence published in 2019 by the GA4GH acknowledges possible re-identification risks and offers possible mitigation strategies (Fiume *et al*., 2019). One strategy is using aggregate beacons. Aggregation process involves querying multiple beacons and joining their responses. A “yes” answer means at least one beacon contains the queried variant; a “no” answer means none has the desired allele. Such an approach leads to having more data points than the individual beacons, which, as also suggested by Shringarpure and Bustamante (2015), makes it harder for the attacker to pinpoint the origin of a “yes” answer. One example of such is the Conglomerate Beacon. However, this strategy also results in a substantial utility loss for the users (researchers) as they might have to apply for access to all individual datasets if they find out that at least one of the beacons have the variant they are interested in. The second suggestion is the usage of participant budgets as suggested by Raisaro *et al*. (2017). This strategy assigns a personal budget to each participant and if many rare SNPs (i.e., relatively more informative and identifying SNPs) of a participant are queried, the algorithm takes them out of the system (i.e., it does not provide a “yes” response if that person is the only carrier of that SNP in the beacon). This seems sensible, yet, in von Thenen *et al*. (2018), authors show that an attacker, by inferring the responses of a beacon via linkage disequilibrium between the SNPs in a population, can get around these budgets. Considering the individual NA12272 from the HapMap project in a beacon of 65 CEU individuals (constructed from the HapMap project), they show that while the Optimal Attack requires 7 queries for re-identification, the QI attack can identify this person with only 5 queries, before the budget expires.

Shringarpure and Bustamante (2015) suggested inclusion of control samples in a beacon. Similar to the aggregate beacon strategy, this decreases the usability and utility of the system since controls, who do not carry the phenotype which the beacon is associated with and who are not relatives of the people in the beacon, would result in flipping of many irrelevant “no” answers to “yes”. In this work, we investigate the feasibility of adding relatives of individuals to a beacon as a countermeasure. Adding relatives still results in a utility loss, however, as shown in Section 3 the loss is not significant given the fact that most SNPs are shared between the victim and their relatives. Moreover, in beacons of heritable diseases, a relative is more likely to be related to the trait than a random control individual. Thus, the utility loss caused by the proposed approach will be less compared to adding random controls. Yet, we show that this creates a major confusion for the attacker. As clearly shown in various settings, the power curves for the state-of-the-art attacks shift right, which indicates that the number of required queries substantially increase to achieve the same re-identification power. In many cases, the power does not even reach to 100%, which means the attacker cannot have high confidence about the success of the attack.

As discussed, in RTF method, Bu *et al*. propose flipping some “yes” responses into “no” after checking a condition for all beacon participants. In our proposed mitigation mechanism, we do the opposite: due to the added relatives on the target individual, our proposed mechanism results in some accuracy loss by flipping some originally “no” responses to “yes”. Thus, we compared our approach with Bu *et al*. in terms of accuracy of beacon responses, especially for queries that are received form rare SNPs. For the comparison, we used the following settings: for Bu *et al*., beacon size is 40 and beacon considers all 40 individuals as potential targets (as also suggested in the original work). In our scheme, original beacon size is 40 and we add both parents of all these 40 individuals (to protect privacy of all beacon participants against the re-identification attacks). Bu *et al*. shows that RTF approach reduces the re-identification power of the attacker to an insignificant level. We also show (in Section 3) that adding both parents of a potential target provides a comparable privacy for beacon participants. Thus, we compared these approaches only based on their utility loss in beacon responses. Our results show that considering responses of the beacon for rare SNPs (for SNPs with MAF value smaller than 0.07), accuracy loss in Bu *et al*. is 25%, while accuracy loss of our proposed mechanism is 16%. Furthermore, considering the main functionality of a genomic data sharing beacon (that it informs a researcher about the existence of a genome in a database), changing “yes” responses into “no” (as in the RTF method) will cause the researcher falsely eliminate the corresponding beacon. However, changing “no” responses into “yes” (as in our proposed technique) will only cause false which will lead to an unnecessary acquisition of the dataset. These results also show that our proposed mitigation mechanism protects beacon participants against re-identification attacks while also preserving the utility of beacon responses.

We also investigated the scenario, in which the attacker also has knowledge about the genomes of the victim’s relatives. For example, we assumed that the attacker has the SNPs of both victim and their mother (and/or father). The initial idea is, if the attacker applies the attacks by using the SNPs that differentiates between relatives and the victim, the power of the attack will reach to 100%. So, the power decrease we achieve by existence of a relative in the beacon will be ineffective. However, in practice, launching the QI attack by using these differentiated SNPs will not be effective since these SNPs are sparse and are less likely to be in linkage disequilibrium. Thus, they are less likely to be correlated to enable inference of the beacon answers To evaluate the power of this new scenario, a new power calculation approach is needed which we will consider as a future work.

We show that the membership inference risk of a person decreases when her relatives are added to the beacon. one might question the risk for the relatives. The membership inference risk for relatives are expexcted to be similar to the child’s since the protection works two ways, symmetrically. That is, the shared SNPs that causes *confusing* “Yes” responses for the attacker protects both the victim and the relative. Then, the risk for both of them should be on par assuming (and likely) they are from the same population.

One drawback of this mitigation strategy is the additional sequencing cost of the relatives. Moreover, the technique depends on relatives giving consent to sharing their data, which also puts them under re-identification risk. However, the protection effect is symmetric for the victim and the relatives. To circumvent these problems, one can opt for simulating relative data, as we did in this work due to the unavailability of large family studies required to quantify the power of re-identification attacks.

## 5 Conclusion

In this paper, we have proposed a mitigation technique against reidentification attacks for genomic data sharing beacons. The existing countermeasures to prevent re-identification attacks in beacons are shown to be ineffective since they either proved to be vulnerable against the attacks or they cause a significant decrease in beacon’s utility. Our proposed technique relies on inheritance and it is based on adding genomes of a victim’s family members to the beacon in order to mitigate the reidentification attacks. We have shown via experiments that adding at least one family member of the victim to the beacon results in a significant decrease in the power of the re-identification attacks. We have also shown the effect of adding different family members to the beacon to the power of the attacker. Furthermore, the proposed technique does not cause a substantial utility loss in beacon’s responses. In particular, we have shown that the utility loss is significantly smaller for SNPs with low MAF values (which are of high importance for the researchers due to their associations with complex diseases).

## Footnotes

1 https://beacon-network.org/

